# eIF3 subunit M regulates blood meal digestion in *Rhodnius prolixus* affecting ecdysis, reproduction and survival

**DOI:** 10.1101/2022.07.16.500302

**Authors:** Pilar Ameijeiras, Natalia Capriotti, Sheila Ons, Pedro L. Oliveira, Marcos Sterkel

## Abstract

In triatomines, blood-feeding triggers many physiological processes, including post-embryonic development and reproduction. Different feeding habits, such as hematophagy, can shape gene functions to meet the challenges of each type of diet. A comparison of transcriptomic and proteomic data indicates that post-transcriptional regulation of gene expression is crucial in triatomines, so we evaluated the impact of RNAi silencing of the eukaryotic translation initiation factors 3 subunit m (eIF3m) in *R. prolixus* physiology. We showed that eIF3m is essential for correct digestion, affecting the processes triggered by a blood meal. The silencing of this gene inhibited moulting and caused the premature death of nymphs, while in adult females inhibited oviposition and increased resistance to starvation. Male survival was not affected by eIF3m knockdown. The information regarding the eIF3m function in insects is scarce. The phenotypes observed in *R. prolixus* upon eIF3m gene silencing are different and more severe than those described in *Drosophila melanogaster*, pointing to the particular importance of this gene in triatomines.

**Summary statement:** The information provided here indicates the importance of mRNA translation in modulating growth, reproduction, lifespan and starvation resistance in triatomine vectors.

## 1. Introduction

Triatomines, as all blood-sucking arthropods, feed on hosts that are thousands of times larger than themselves, so the timing of feeding represents a high risk to their survival. The sanitary relevance of this insect subfamily is because of their role as vectors of *Trypanosoma cruzi*, the parasite that causes Chagas disease. Despite hematophagy arose on many independent occasions during the evolution of arthropods (Lehane, 2005), a common adaptive strategy has been to decrease the frequency of feeding, compensating with the ingestion of enormous volumes of blood equivalent to several times their body weight during each visit to the host (Lehane, 2005). The adaptations that allow blood-sucking arthropods to tolerate their diet constitute a unique operating model regarding cell signalling, immunity, and metabolism (Sterkel et al., 2017). A blood meal impacts triatomine physiology and triggers several processes, such as diuresis, digestion, moulting in nymphs and reproduction in adults, involving a massive synthesis of new proteins synchronized with digestion (Guarneri and Lorenzo, 2021).

According to the so-called “ribosome filter” hypothesis, eukaryotic translation initiation factors (eIF) recruit specific sets of mRNAs on the ribosome without causing a drastic change in the existing mRNA pool (Mauro and Edelman, 2007). eIF3 is required to form a stable 40 S preinitiation complex and stabilizes the binding of the eIF2-GTP-Met-RNAt complex to the 40S ribosomal subunit (Chaudhuri et al., 1999). eIF3 contains 13 subunits, named alphabetically as eIF3a-eIF3m (Zhou et al., 2008). This complex has constituent subunits conserved in all eukaryotes (eIF3a, eIF3b and eIF3c, eIF3g and eIF3i) and variable subunits (Masutani et al., 2007). eIF3 complex associates with the mRNA 5’ untranslated portion of mRNAs related to cell growth, cell cycling, differentiation and apoptosis (Lee et al., 2015). It was proposed that eIF3m and eIF3e define two different eIF3 complexes with a distinct affinity for the different mRNAs, thus selecting the set of mRNAs to be translated and constituting a post-transcriptional mode of regulation of gene expression (Zhou et al., 2005).

In humans, eIF3m regulates the expression of tumorigenesis-related genes in colon cancer (Goh et al., 2011) and promotes the malignant phenotype of lung adenocarcinoma by the up-regulation of oncogene *CAPRIN1* (Liu et al., 2021). eIF3m silencing inhibits herpes virus protein translation but has little effect on cellular RNA or protein expression and is not cytotoxic (Cheshenko et al., 2010). In mice, eIF3m eIF3m is critical for eIF3 structure and is required for embryonic development, homeostasis, and organ size control (Zeng et al., 2013). The knockdown of eIF3m has a limited impact on mRNA-specific translation but affects ribosome biogenesis and transcription in mice’
ss liver (Smekalova et al., 2020).

In insects, the physiological role of eIF3m has been studied in *D. melanogaster*, in which eIF3m (*Tango7*. CG8309) is a pro-apoptotic factor (Chew et al., 2009). It positively regulates *Dronc* caspase levels through a post-transcriptional mechanism. Its silencing prevents death induced by multiple caspase-dependent apoptotic stimuli (Chew et al., 2009). However, eIF3m has functions in cells that are not destined to die, such as cell remodelling by the apoptosome (D’Brot et al., 2013). eIF3m regulates the caspase-dependent remodelling process necessary for sperm development (D’Brot et al., 2013) and regulates *Dronc* caspase activity at the cortex in salivary glands of *D. melanogaster* (Kang et al., 2017).

The eIF3m subunit is conserved from fission yeast to higher eukaryotes, but it is absent in budding yeast (Zeng et al., 2013), and it is the only unidentified eIF3 subunit in trypanosomatid genomes (Rezende et al., 2014). After sequencing the *R. prolixus* genome, the first triatomine genome available (Mesquita et al., 2015), several transcriptomic studies appeared correlating mRNA level profiles of different tissues with physiology, immunity, reproduction and development (Coelho et al., 2021; Latorre-Estivalis et al., 2020, 2017; Leyria et al., 2020a, 2020b; Ribeiro et al., 2014). However, the results obtained with a proteomic approach in the midgut showed an expression profile distinct from the mRNA, suggesting that post-transcriptional regulation of gene expression is crucial (Ouali et al., 2021). These observations led us to evaluate the impact of RNAi silencing of eIF3m expression in *R. prolixus* physiology. Its knockdown reduced the digestion rate affecting development, reproduction and survival. Interestingly, the phenotypes observed upon silencing the expression of eIF3m in *R. prolixus* are different and more severe than those reported in *D. melanogaster*, in which the effects of its silencing were mild and tissue-specific (Chew et al., 2009; D’Brot et al., 2013; Kang et al., 2017). The results point to a particular and crucial role of eIF3m in triatomine physiology.

## 2. Materials and methods

### 2.1. Rearing of insects

Experiments were performed with animals obtained from a colony in the Instituto de Bioquímica Médica Leopoldo de Meis, Universidade Federal do Rio de Janeiro (UFRJ), Brazil, and in the Centro Regional de Estudios Genómicos, Universidad Nacional de La Plata (CREG), Argentina. In both cases, insects were maintained at 28 °C and 50– 60% relative humidity under a photoperiod of 12 h of light/12 h of dark. In UFRJ, the insects were fed on rabbit blood at 3-week intervals. In CREG, insects were fed on chickens at 3-4 weeks intervals. All the animal work was conducted according to the guidelines of the institutional care and use committee (Committee for Evaluation of animal Use for Research from the Federal University of Rio de Janeiro), which is based on the National Institutes of Health Guide for the Care and Use of Laboratory Animals (ISBN 0-309-05377-3). The protocols received registry number 149-19 from the Animal Ethics Committee (Comissão de Ética no Uso de Animais, CEUA). Technicians at the animal facility at the Institute of Medical Biochemistry (UFRJ) performed all aspects related to rabbit husbandry under strict guidelines to ensure careful and consistent handling of the animals. Biosecurity considerations are in agreement with CONICET resolution 1619/2008, which is in concordance with the WHO Biosecurity Handbook (ISBN 92 4 354 6503)

### 2.2. RNA isolation and complementary DNA (cDNA) synthesis

*R. prolixus* tissues were dissected in ice-cold PBS (NaCl 0.15 M, Na^+^-phosphate 10 mM, pH 7.4). The total RNA from different tissues was extracted using TRIzol reagent (Ambion, USA), according to the manufacturer’s instructions. Following treatment with DNAse I (Invitrogen, USA), first-strand cDNA synthesis was performed using 1 µg total RNA with MMLV Reverse Transcription Kit (Applied Biosystems, Brazil) and poly-T primer, according to the manufacturer instructions.

### 2.3. Synthesis of double-stranded RNA (dsRNA)

Two pairs of specific primers for the eIF3m (RPRC009587) gene containing the T7 RNA polymerase binding sequence at 5’ end, required for dsRNA synthesis, were designed. We ruled out off-target effects by injecting two dsRNA covering different regions of the eIF3m mRNA in independent experiments. This was done to confirm that the phenotypes observed were due to eIF3m knockdown and not to the possible unspecific silencing of other genes due to sequence similarity. dsRNA-1 was designed in exons 1 and 2 (primers: Fw 5’
s-TAATACGACTCACTATAGGGAGATCTGGACGTTCCGGCAGT-3′ and Rv 5’-TAATACGACTCACTATAGGGAGATGGGCATCTTCCCTGGC-3′). dsRNA-2 was design in exon 3 (primers: Fw 5′-TAATACGACTCACTATAGGGAGAGCCCTAGCCGATCCTAACACT-3’and Rv 5’-TAATACGACTCACTATAGGGAGATTTCATTAGCACCTTCAGCC-3’)

A fragment from the β-lactamase gene, absent in the *R. prolixus* genome, was PCR-amplified from the pBluescript plasmid (primers: Fw 5-TAATACGACTCACTATAGGGGAACTGGATCTCAACAG-3’
s and RV: 5′-TAATACGACTCACTATAGGGGGATCTTCACCTAGATC-3′) and used as a control to assess the putative unspecific effects of dsRNA injections. All the PCR products were sequenced (Macrogen, Korea) to confirm their identity. One µl of the PCR product was used for a second PCR using T7-full promoter primer (5′-ATAGAATTCTCTCTAGAAGCTTAATACGACTCACTATAGGG-3′). The product of the second PCR was used as the template for dsRNA synthesis. The dsRNAs were in vitro transcribed using T7-RNA polymerase (Invitrogen, USA), according to the manufacturer’s instructions. dsRNAs were precipitated with isopropanol and resuspended in ultrapure water (Milli-Q). The dsRNAs were visualized by agarose gel (1% w/v) electrophoresis to verify size, integrity and purity. The dsRNAs were quantified from images of the gels using the software ImageJ (National Institute of Health, USA). The dsRNAs were stored at −20°C until use.

### 2.4. RNAi to determine loss-of-function phenotypes

Fourth instar nymphs, adult males and females *R. prolixus* were injected into the thorax with 2 μg of each dsRNA dissolved in 2 μl of ultrapure water using a 10 μl Hamilton microsyringe. Control insects were injected with 2 μg of β-lac dsRNA. Insects were fed seven days after the dsRNA injection, which was considered day 0 PBM. On that day, the intestine of some starved insects was collected in Trizol reagent (Invitrogen, USA) to check the efficacy of gene knockdown by qPCR. Mated males and females that had been fed previously once during the adult stage were used to perform the experiments. Adult insects were injected 21 days after feeding, while fourth instar nymphs were injected 15-21 days after moulting.

### 2.5. Quantitative polymerase chain reaction (qPCR)

Total RNA was extracted from the intestine (anterior midgut, posterior midgut and rectum) of females, males and fourth instar nymphs seven days after dsRNA injection. The cDNAs were synthesised as previously described. Specific primers targeting the eIF3m gene (Fw 5’-GGAAGAAGTAGAGGCTTTCGTG-3’ and Rv 5’-TGTTCCCATTGGGCTACTCC-3’) were designed to amplify a different region from that amplified by the RNAi primer pairs to prevent dsRNA amplification that may be retrotranscribed during synthesis of the cDNA together with insect RNA. They were also designed in different exons (exons 3 and 4) to prevent genomic DNA amplification. Primers were tested for dimerization, efficiency (85.7%) and amplification of a single product. The Glucosa-6-fosfato dehydrogenase (G6DPH) gene was used as references (housekeeping) (Fw 5’-AGCCTGGAGAAGCGGTTTACGTTA-3’ and Rv 5’-GTGAGCCACAGAATACGTCGAGT −3’) (Omondi et al., 2015). qPCR was performed using Master Mix qPCR Sybr green (Productos Bio-logicos, Argentina) under the following conditions: 95 °C for 5 minutes, followed by 40 cycles of 95 °C for 30 seconds, 60 °C for 30 seconds and 72 °C for 15 seconds. Finally, the melting temperature of the PCR product generated was calculated (1 cycle of 95 °C for 30 seconds, 65 °C for 30 seconds and 95 °C for 30 seconds) in an Agilent AriaMx instrument. For each sample, ΔCT values were calculated (CT eIF3m - CT G6DPH). The 2e^-ΔCT^ values obtained for dseIF3m and dsβ-lac-injected insects were used to evaluate gene-silencing efficacy (Livak and Schmittgen, 2001).

### 2.6. Haemoglobin quantification

The anterior midguts were collected immediately after the blood meal (between 0.5–2 h), 2, 4, 7 and 14 days PBM and homogenized in 250 µL of PBS (10 mM Na– phosphate, 0.15 M NaCl, pH 7.4) in a 1.5 ml plastic tube. The volume was adjusted to 1 ml with PBS. Haemoglobin content was assayed with a colorimetric kit (K023 kit, Bioclin, Brazil). Two µl of each sample were mixed with 198 µl of reagent for measurements. The standard curve was performed with the provided standard (15 mg/ml). 5, 4, 3, 2, and 1 µl of the standard were mixed with 195, 196, 197, 198 and 199 µl of reagent, respectively. The absorbance of the supernatants was read at 540 nm with a plate spectrophotometer (Spectramax M3, Molecular Devices, United States).

### 2.7. Protein content measurement in ovaries

The ovaries from the same females dissected for haemoglobin quantification were collected in 300 µl of PBS and macerated in a 1.5 ml plastic tube. The volume was adjusted to 1 ml with PBS. 50 µl were used for protein quantification by the Bradford method, according to the protocols described by (Bradford, 1976). Bovine serum albumin (BSA) was used as a standard for protein quantification.

### 2.8. Oviposition and eclosion

Fully engorged females were individually separated into vials and kept at 28 °C and 50–60 % relative humidity under a photoperiod of 12 h of light/12 h of dark. The number of eggs laid by each female was counted daily. The eclosion ratios were calculated by dividing the number of hatched first instar nymphs by the number of eggs laid by each female.

### 2.9. Survival and ecdysis measurements

Insect survival and/or ecdysis occurrence was scored daily after the blood meal (PBM).

### 2.10. Statistical Analysis

At least two independent replicates were performed for each experiment. Each replicate included N=7–20 insects per experimental group. The data from different replicates were combined into a single graph for the design of the figures. The statistics were performed on both, each independent experimental replicate and the final data containing the information from the different replicates combined. The p values indicated in the text are from the data of the independent experiments combined. Statistical analysis and graphs designs were performed using Prism 8.0 software (GraphPad Software). Survival was evaluated by Kaplan-Meier curves and Log-rank (Mantel-Cox) test. Statistical differences between the experimental and control groups in oviposition, egg hatching and qPCR were evaluated by unpaired t-test. Time course differences between groups in haemoglobin and protein content, and in ecdysis were assessed by Two-way ANOVA.

## 3. Results

A high reduction of eIF3m mRNA levels was achieved in nymphs and adult insects injected with dsRNA (Fig. S1A-C). Even though similar volumes of blood were ingested during feeding, the amount of haemoglobin in the anterior midgut was higher in eIF3m silenced insects by day 7 PBM (Two-way ANOVA. p<0.0001; N=9), indicating that eIF3m knockdown reduced blood meal digestion rate. By day 14 PBM, control females had already digested all the blood meal, while eIF3m-silenced insects still possessed ∼30% of the ingested haemoglobin (Fig. 1A). Coincident with the reduction in the rate of blood meal digestion, the blockage of ovary development was observed. The protein content of the ovaries was drastically reduced on days 7 and 14 PBM (Two-way ANOVA. p<0.01; N=6) compared with controls, while no differences were observed on days 2 PBM (before oviposition begins) or 21 PBM (after oviposition finished) (Fig. 1B). In consequence, the oviposition was drastically reduced (Fig. 2A-B). The few eggs that were laid presented malformations (Fig. S2), and any nymph hatched from them (Fig. 2C-D). Therefore, reproduction was completely blocked.

**Figure 1:**
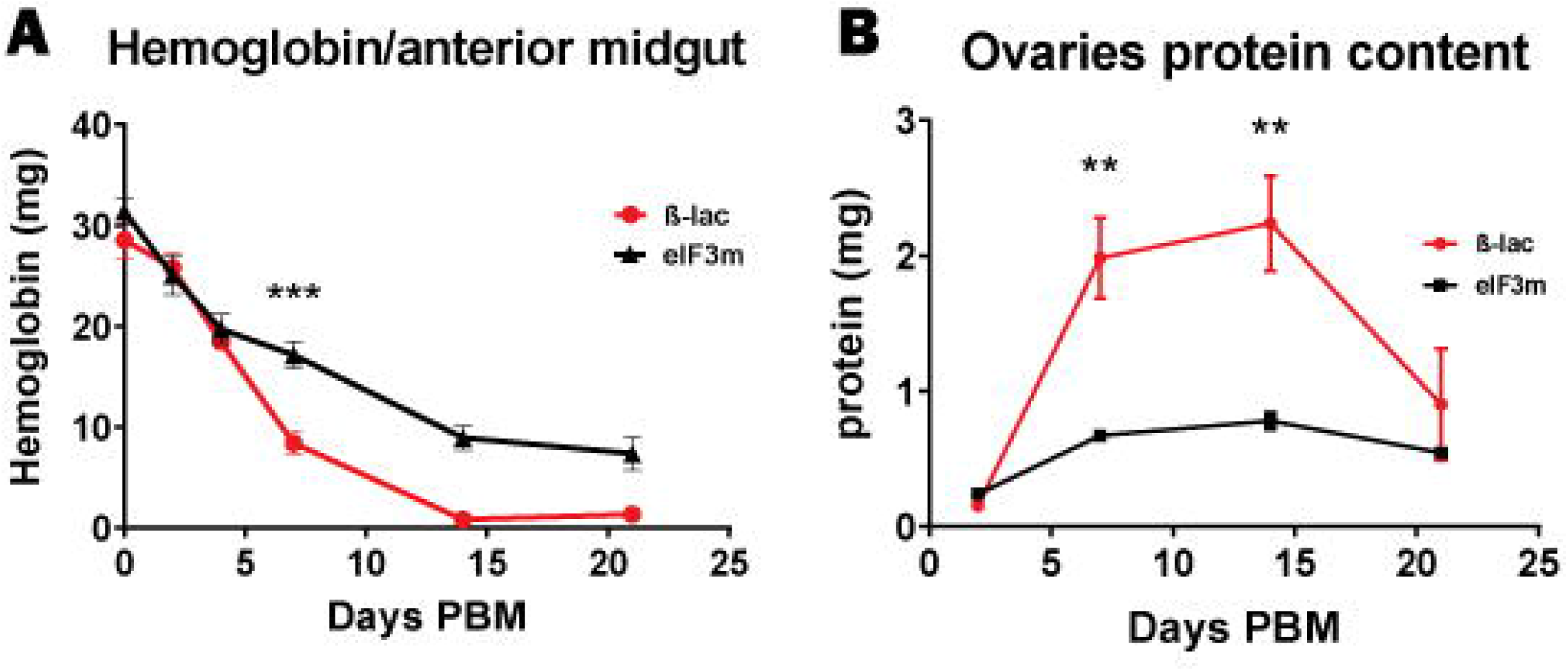
Silencing of eIF3m reduces digestion rate and prevents ovaries development. **A:** Amount of haemoglobin in the anterior midgut. **B:** Quantity of protein in the ovaries. Two-way ANNOVA was performed on the different time points to evaluate the difference between the control and dseIF3m injected group. ***p<0.001, **p<0.01. Two independent experiments were carried out, each with N=3-5 insects per experimental group in each time point. Data from both experiments were combined into a single graph. Data are plotted as mean + s.e.m.

**Figure 2:**
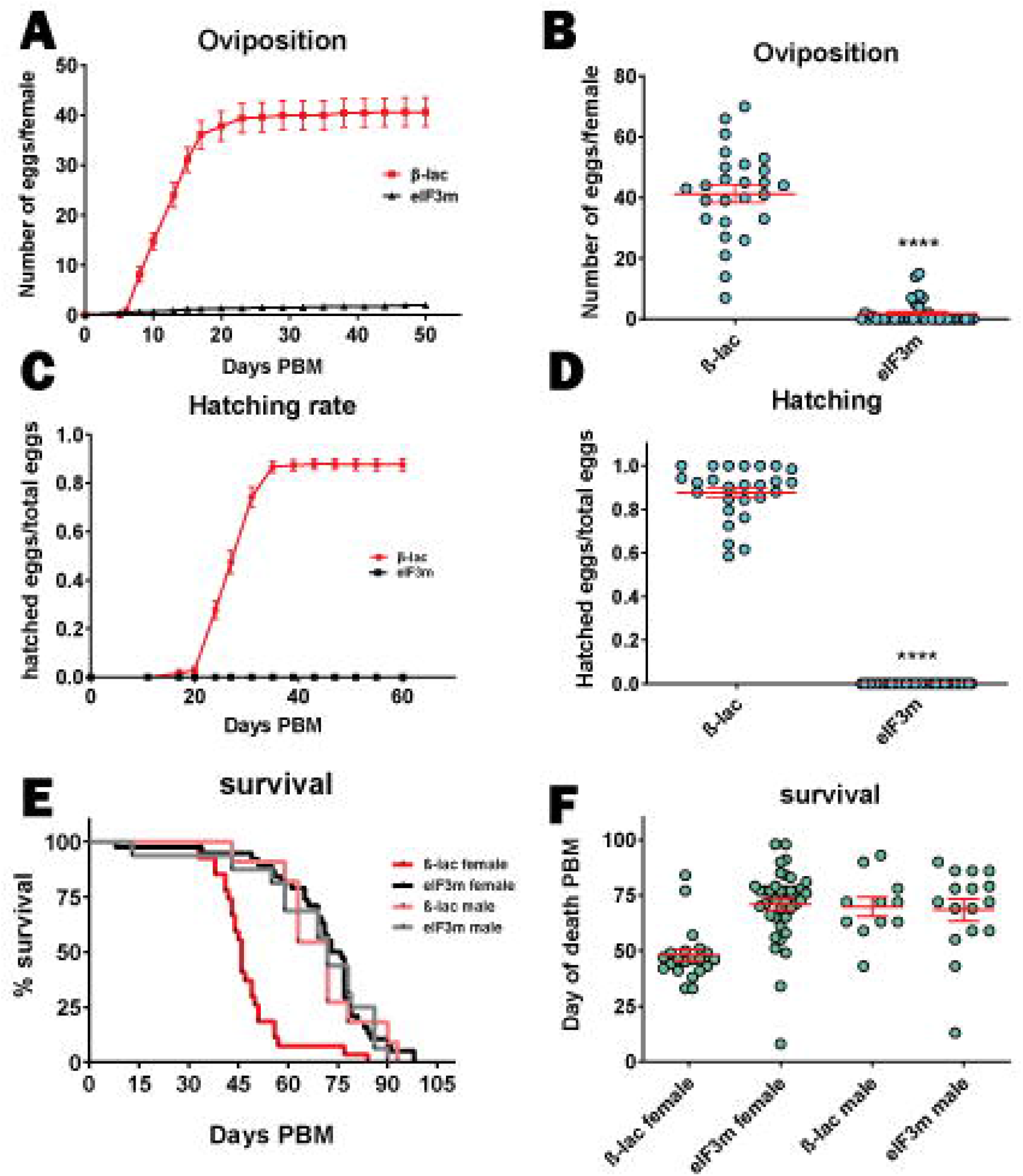
eIF3m silencing impairs reproduction and increases resistance to starvation in females. **A-B:** Oviposition. **C-D:** The hatching of eggs. An unpaired t-test was performed. ****p<0.0001. Data are plotted as mean + s.e.m. **E-F:** Female and male survival. Data are plotted as Kaplan-Meier curves (E) and as the death day post-blood meal (F). Control females presented a reduced survival compared to the other groups (Log-rank (Mantel-Cox) test. P<0.0001). Four independent experiments were carried out with N=5-12 insects per experimental group. Data from independent experiments were combined into a single graph.

eIF3m-silenced females were more resistant to starvation and survived longer than controls after a blood meal (Fig 2E-F). Control female survival was 46 +/- 2.6 (Mean+/- S.E.M) days PBM, while eIF3m-silenced females lived for 70.97 +/- 2.7 days (p<0.0001). In contrast, a similar survival period was observed in males upon eIF3m silencing. Both control (69.82+/-4.26 days PBM) and eIF3m-silenced males (68.38+/-4.91 days PBM) survived for a period comparable to eIF3m-silenced females (Fig 2E-F). These results suggest that the knockdown of eIF3m caused a switch towards survival at the expense of reproduction in the females. This reallocation of resources made the females more resistant to starvation, and they survived as long as the males. In fourth instar nymphs, the knockdown of eIF3m prevented ecdysis (Two-way ANOVA P<0.0001; N=47) and reduced survival (P<0.0001). The nymphs did not moult despite their survival being longer than the expected ecdysis period (Fig 3A-B).

**Figure 3:**
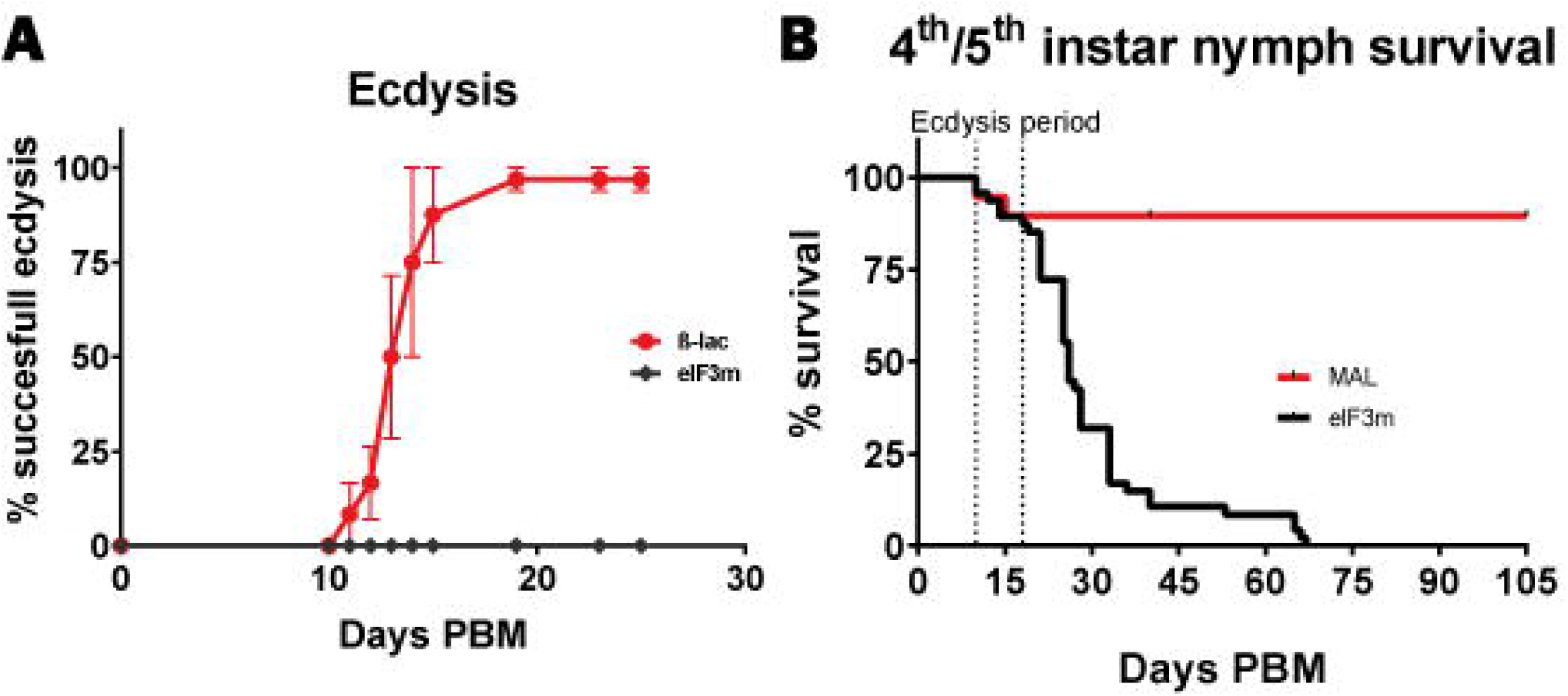
eIF3m silencing prevented ecdysis and reduced nymphs’ survival. **A: Ecdysis**. Two ways ANOVA was performed (P<0.0001). Data are plotted as mean + s.e.m. **B: Survival**. Data are plotted as Kaplan-Meier curves (Log-rank (Mantel-Cox) test. P<0.0001)). Three independent experiments were carried out, each with N=8-15 insects per experimental group. Data were combined into a single graph.

## 4. Discussion

The different feeding habits can shape the function of genes; the loss of function of these genes could have different, sometimes opposite, effects in related species with different diets. For example, inhibition of the tyrosine catabolism pathway extends the lifespan of *D. melanogaster* (Parkhitko et al., 2020) and *Caenorhabditis elegans* (Ferguson et al., 2013; Lee et al., 2003) but causes hematophagous arthropods death after a blood meal due to tyrosine accumulation and precipitation (Ramirez et al., 2021; Sterkel et al., 2021, 2016; Sterkel and Oliveira, 2017). Here we demonstrated that the silencing of the *eIF3m* gene, different from the phenotypes described in *D. melanogaster* that were only mild and tissue-specific (D’Brot et al., 2013; Kang et al., 2017), drastically affected *R. prolixus* physiology, causing a delay in blood meal digestion that affected moulting, reproduction and survival.

In nymphs, the knockdown of eIF3m prevented ecdysis and reduced survival. The insects did not moult despite their survival being longer than the expected moulting period, indicating that the reduced survival was not due to problems during the ecdysis process. If it were the case, the nymph should die during the expected ecdysis period, as observed for Crustacean Cardioactive Peptide (Lee et al., 2013), Orcokinin A (Wulff et al., 2017) and Ecdysis Triggering Hormone knockdowns (Sterkel et al., 2022), presenting the characteristic phenotype of interrupted ecdysis (Wulff et al., 2017).

The phenotypes observed in females upon eIF3m silencing are similar to those reported in non-hematophagous organisms upon dietary restriction (DR). DR extends lifespan while reducing the reproductive fitness of females in many animal species (Green et al., 2022; Moatt et al., 2020), except for *Musca domestica* (Cooper et al., 2004). Lifespan extension by DR has been proposed to be due to a shift in resources from growth and reproduction towards somatic maintenance allowing the animals to survive nutrient-poor environments until they find favourable conditions to reproduce (Attisano et al., 2012; Moore and Attisano, 2011). Low levels of methionine and its derivate metabolite S-adenosyl-methionine (SAM) were indicated to be involved in the phenotypes observed upon DR (Green et al., 2022; Obata et al., 2018). The depletion of dietary methionine (or inhibition of SAM synthesis) reduces the division rate of intestinal stem cells. Homeostatic epithelial turnover is suppressed when SAM is lost (Obata et al., 2018). eIF3m knockdown may emulate DR by reducing digestion rate and limiting nutrient availability.

TOR (target of rapamycin) kinase is a nutrient and amino acid sensor that plays a central role in mediating the lifespan-extending effect of DR (Papadopoli et al., 2019). Inhibiting the TOR pathway extends lifespan and suppresses protein translation, stimulates autophagy, and promotes metabolic health (Papadopoli et al., 2019). Besides, it was identified a protein sensor for SAM, named SAMTOR, that controls TOR complex 1 (TORC1) activation (Gu et al., 2017). In *R. prolixus*, amino acids from blood digestion trigger the downregulation of superoxide via the TORC pathway in the midgut (Gandara et al., 2016). Therefore, a plausible hypothesis is that eIF3m silencing might, by reducing digestion rate, reduce TOR complex signalling.

Sex differences in terms of lifespan and the costs of reproduction remain poorly understood in animals. In *D. melanogaster*, the magnitude of the response and the food concentration that minimized adult mortality differed significantly between the sexes. Female flies subject to DR lived up to 60% longer than fully-fed females, whereas males lived only up to 30% longer (Magwere et al., 2004). In *R. prolixus*, we did not observe differences in survival between eIF3m-silenced and control males. This fact also reinforces the hypothesis that the extended survival observed in eIF3m-silenced females was due to resource reallocation from reproduction to somatic maintenance, making them as resistant as males to starvation.

Digestion of a blood meal represents a challenge for intestine homeostasis due to the release of high and potentially toxic quantities of haem, iron, amino acids and a dramatic increase in the microbiota (Eichler and Schaub, 2002; Sterkel et al., 2017). The damaged or senescent enterocytes must be continually replaced by new cells to maintain epithelial integrity. Since in *D. melanogaster* eIF3m is a pro-apoptotic factor (Chew et al., 2009), eIF3m knockdown may prevent enterocyte apoptosis and the cellular turnover of the intestine, affecting its functions (such as the digestion of blood meal). Further experiments are required to test these hypotheses.

Many long-lived mutants with reduced reproduction were described (Shmookler Reis et al., 2009). Reproduction tends to shorten lifespan in most organisms however, in some cases, the trade-off between reproduction and lifespan can be decoupled (Flatt, 2011). In *D. melanogaster*, mutations in the insulin receptor gene extend lifespan but females are infertile with no-vitellogenic ovaries (Tatar et al., 2001). In *R. prolixus*, insulin receptor deficiency improved longevity and reduced triacylglycerol storage in the fat body, whereas blood digestion remained unchanged for seven days after a blood meal. The females exhibited smaller ovaries and a marked reduction in oviposition (Silva-Oliveira et al., 2021). The knockdown of RpNOX5 and RpXDH prevents blood digestion, impairs egg production and induces early mortality in *R. prolixus* (Gandara et al., 2021). Besides, blood digestion relies on the massive synthesis of proteins, such as digestive enzymes and perimicrovilar membrane proteins. Since eIF3m is involved in translation, it is necessary to investigate how eIF3m silencing affects gene expression. For this, transcriptomic and quantitative proteomics experiments must be performed.

Similar to the phenotypes we observed in *R. prolixus* upon eIF3m knockdown, in C. *elegans*, the knockdown of eukaryotic translation initiation factor 4 gamma 3 (eIF4G3) and S6 kinase, which resulted in the differential translation of genes, slows development, reduces fecundity and increases resistance to starvation and lifespan (Pan et al., 2007). Altogether, these findings indicate a pleiotropic effect of mRNA translation in modulating growth, reproduction, lifespan and the ageing process in different animals. Further studies are required to address the molecular mechanisms underlying the phenotypes observed upon eIF3m silencing in *R. prolixus*. Besides, the function of this gene must be studied in other hematophagous arthropods to evaluate if its physiological importance is conserved among blood-feeders. A deeper understanding of the molecular mechanisms underlying the phenotypes observed upon eIF3m silencing may facilitate the identification of new targets for vector control.

## 5. Acknowledgments

We thank Raúl Stariolo from Centro de Referencia de Vectores for their generous assistance in insect rearing. We also thank Rolando Rivera Pomar (CENDIE-UNOVA) and his lab team for providing the microscope and assistance to perform experiments related to this manuscript.

## 8. Author contributions

P.A. Performed the experiments and analysed the data. N.C. Performed the experiments and contributed critical discussion of the experiments and the data. S.O. Designed the experiments and wrote the paper. P. L. O. Designed the experiments and wrote the manuscript. M. S. designed and performed the experiments, analysed the data and wrote the manuscript. All of the authors discussed the results and read and contributed to the final version of the manuscript.

